# Multifaceted integration - memory for faces is subserved by widespread connections between visual, memory, social and auditory networks

**DOI:** 10.1101/382739

**Authors:** Michal Ramot, Catherine Walsh, Alex Martin

## Abstract

Face memory abilities are at the core of human social interaction, yet these vary widely within the general population, ranging from developmental prosopagnosia to “super-recognizers”. Previous work has focused mainly on the contribution of the well described face network to this variance. However, given the nature of the face memory task, and the social context in which it takes place, we were interested in exploring how the collaboration between different networks outside the face network (measured through resting state connectivity) affects performance on face memory tasks. We discovered that face recognition memory is supported by a wide network of connections between the face patches, memory regions, auditory regions and social networks. Moreover, this network was selective for memory for faces, and did not predict memory for other visual objects, such as cars.

## Introduction

Memory for faces is one of the core capacities of the human mind. The ability to recognize people, and to know whether the person in front of us is familiar or not, is fundamental to our social functioning, a cornerstone of humanity. And yet, within the general, healthy population there is a great degree of variance in terms of ability to remember and recognize both familiar and novel faces ^1-3^.

The neural underpinnings of visual face processing have been studied extensively ^4-8^, resulting in the identification of several patches in the ventral visual stream along the fusiform gyrus such as the occipital face area (OFA), the fusiform face area (FFA), and more recently, a more anterior patch in the ventral anterior temporal lobe (ATL) which are highly selective for face stimuli, responding more to faces than to any other visual category ^9, 10^. These regions, along with the amygdala, parts of the lateral occipital sulcus (LOS) and the posterior superior temporal sulcus (pSTS) are largely regarded as the elements comprising the face network ^11, 12^.

Lesion and stimulation studies have shown that face perception and recognition are impaired when these regions are compromised ^11, 13^. The degree of activation and selectivity within this face network in response to face stimuli has been linked to better performance on face memory tasks in numerous studies ^1, 14, 15^, with patients with congenital prosopagnosia demonstrating significantly reduced selectivity within these regions ^16, 17^. Connectivity between nodes of the face network has been shown to correlate with increased selectivity ^18^, and activity in this region can be used to decode faces ^19^. However, studies investigating the link between resting state functional connectivity and face recognition memory abilities have been scarce and limited in scope, focusing almost exclusively on correlations within the face network itself, and of the face network with early visual cortex. These have shown decreased connectivity between these nodes in individuals with congenital prosopagnosia ^9, 20^, and predictive value for the correlation between OFA and FFA for performance on face recognition tasks ^21^. Studies using Diffusion Tensor Imaging (DTI) have identified reduced structural connectivity along the ventral occipito-temporal cortex, in white matter tracts projecting from the occipital face regions to anterior temporal regions ^22^.

The story of face recognition memory, across the full range of abilities, from cases where it is severely disrupted without any apparent brain insult, such as in congenital prosopagnosia, through typical individuals to those with exceptional abilities, has so far been examined primarily if not exclusively through the face network. Yet surely, given the nature of the processes involved in face recognition and the context in which it is performed, both memory and social networks must also be profoundly involved. We therefore set out on a whole brain search to characterize the networks outside the face patches that underpin face recognition memory, by searching for regions whose connectivity with the ventral face patches predicts face memory ability.

## Results

### Behavioral tests and ROI localization

Prior to the fMRI scan, participants came in for a behavioral testing session, which included administration of the Cambridge Face Memory Test (CFMT) ^23^, a widely utilized measure of face memory ability, as well as the Cambridge Car Memory Test (CCMT) ^24^ in counterbalanced order, outside the MRI scanner. Performance on the two tests was significantly correlated across our 50 participants (r = 0.45 p < 0.0005, two tailed t-test), and they were well matched for difficulty, with no significant difference in the mean scores of the tests (mean score = 78.2 for the CFMT, 75.1 for the CCMT). Participants then went on to do an fMRI scan, which included two rest scans, two face/scene localizer runs, and one movie run. All runs were approximately nine minutes long (see methods for more details). We began by using the face/scene localizer runs to identify the ventral face patches (bilateral OFA, bilateral FFA, and right ATL), as well as bilateral amygdala in each individual participant (N=50). Left ATL was difficult to localize in some participants (congruent with the known right bias for the face network and specifically for ATL ^11^) and was therefore excluded. These regions of interest (ROIs) were defined as 6mm radius spheres around the center of mass of each cluster. Figure 1 shows the location of these regions for a representative participant.

**Figure 1.**
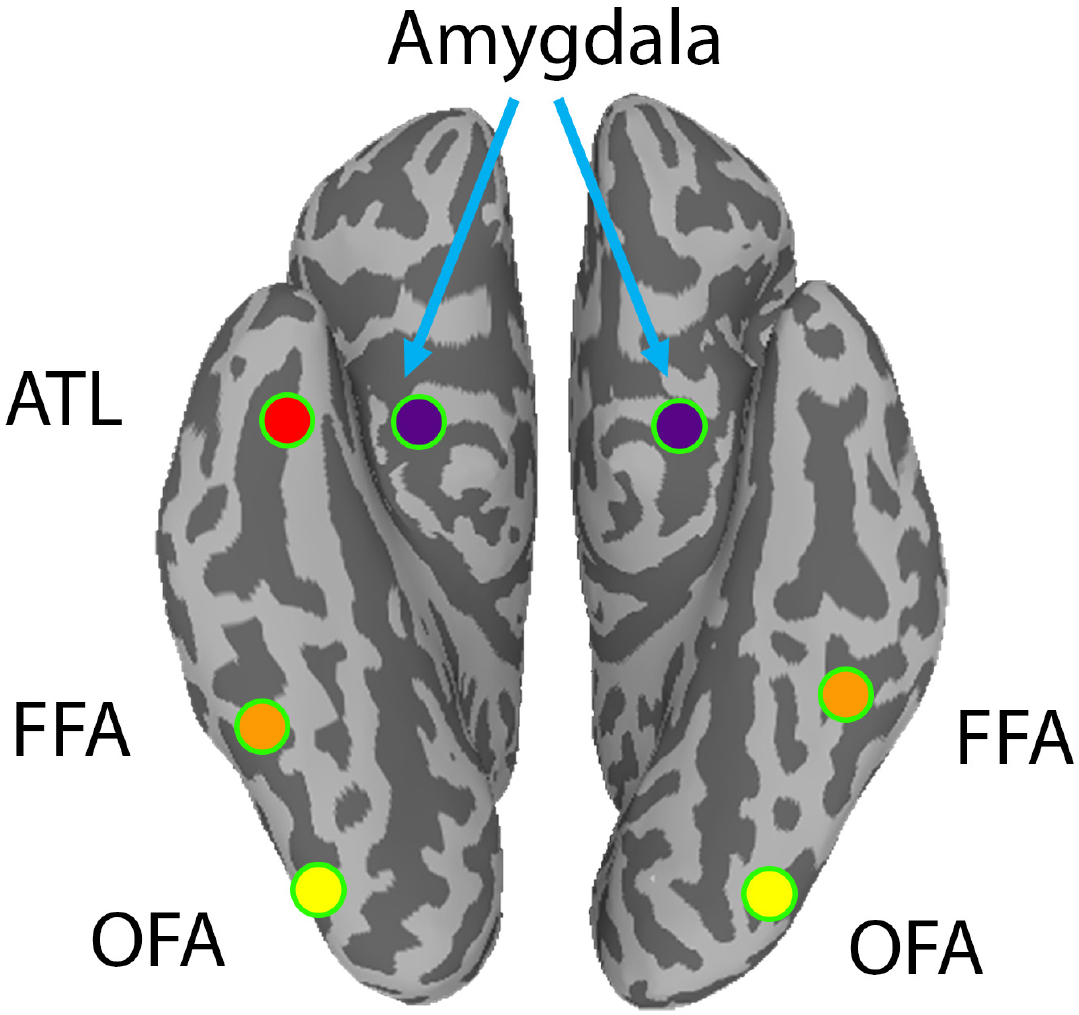
Location of face ROIs. Map showing sample location of the seven individually localized face ROIs, in one representative participant. ATL-anterior temporal lobe face region, FFA–fusiform face area, OFA – occipital face area.

### Face selectivity

Next, we sought to reproduce previous findings linking selectivity for faces within the face regions to performance on the face memory task ^1, 14, 15^. We first defined selectivity as the difference of the beta values of the face > scene condition in the localizer runs for each participant, averaged across the voxels in each of our individually defined face patches. We then correlated this selectivity value across participants, with their performance on the CFMT and CCMT. Selectivity of the right FFA was significantly correlated with performance on the CFMT (r = 0.33, p = 0.026), but not with performance on the CCMT (r = 0.15 p = 0.3). Selectivity of right OFA was trending toward significant correlation with the CFMT (r = 0.28, p = 0.06) but not with the CCMT (r = 0.07, p = 0.62). The other face ROIs were not significantly correlated to either CFMT or CCMT performance. The correlation between selectivity and performance on the CFMT increased when the selectivity was averaged across several of the face ROIs and was strongest when averaged across the three right ventral face patches, right OFA, right FFA and right ATL, as displayed in Figure 2 (r = 0.4, p = 0.007).

**Figure 2.**
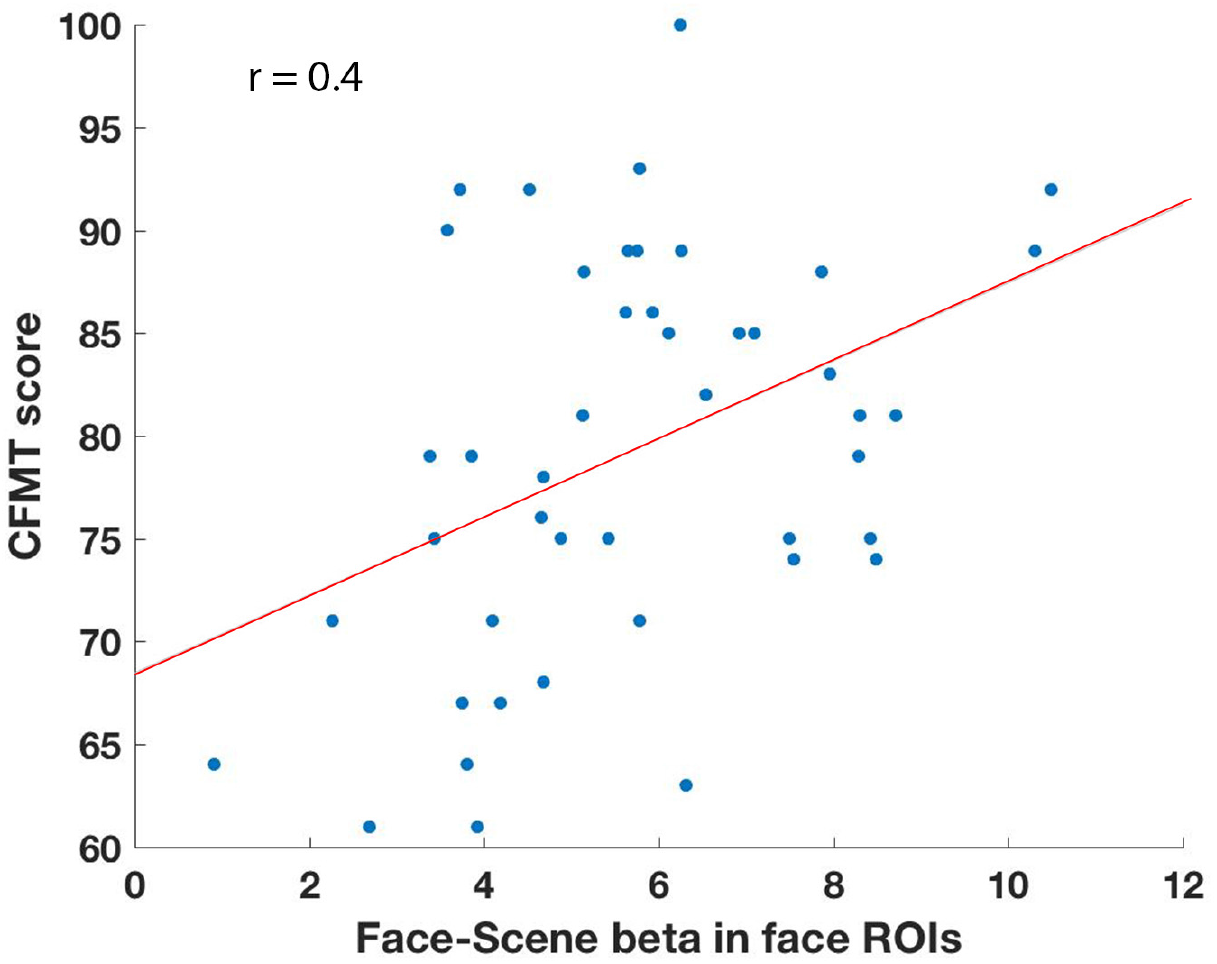
Face selectivity. Face-scene beta during the two localizer runs averaged across all voxels in the three right ventral face patches (right OFA, right FFA and right ATL) shown on the x-axis per participant, with CFMT scores shown on the y-axis. R = 0.4, P = 0.007, N=46, as four participants were excluded because they had been given a different version of the localizer task (see methods).

### Extending the network

Having reproduced the findings relating to the correlation between selectivity and face memory abilities within the face network in the localizer data, we turned to the resting state data. The goal was to search for regions beyond the face network, which interact with the face network in a meaningful way in relation to face memory abilities. To this end, we took each one of our face ROIs as a seed and calculated the correlation of the time course of each voxel in the brain with the time course of that seed during rest, for each participant. We then calculated for each voxel, across participants, the second order correlations with the CFMT score, by correlating for each voxel its correlations to the seed with the CFMT scores. This gave us a measure for each voxel of how predictive its correlation with the face seed region was to behavior, as measured by the CFMT score. We repeated this analysis for each of the 7 face ROIs, and for both rest scans. To validate the findings, we constrained our results for each seed to voxels that had significant second order correlations to behavior across both rest scans, corrected for multiple comparisons using a strict cluster size permutation test (see methods). The corrected maps showing the voxels whose correlation with right ATL, FFA and OFA were significantly predictive of behavior are displayed in Supplementary Figures 1-3, respectively, and are largely overlapping, as were maps for the other seed regions. A similar analysis was conducted with correlation to non-face memory scores, as measured by the CCMT, but no significant clusters were found.

We next identified the peaks of the clusters that were predictive of CFMT scores across the two rest scans and survived correction for multiple comparisons in any of the second order correlation seed maps, and defined those as new ROIs, with 6mm radius spheres. 25 such ROIs were identified, some in medial parietal and medial temporal regions, others in inferior frontal gyrus (IFG), somatosensory regions, along the insula, auditory cortex in the superior temporal gyrus, and the dorsal attention stream. This analysis also picked out a region in the lateral occipital sulcus (LOS) and another in posterior superior temporal sulcus (pSTS), both of which are often included in the face network. These ROIs are shown in Figure 3. This approach was motivated by the hypothesis that it is the connectivity between the face network and other memory/social networks which underlies face memory, congruent with the ROIs identified in this manner. However, we also took a completely data driven approach, ignoring the face ROIs defined by the localizer scans. Instead, we calculated for each participant for each voxel the global connectivity of that voxel (i.e. the average correlation of that voxel to all other voxels), and then calculated the second order correlation of the global connectivity with the CFMT scores. Figure 4 shows the corrected map of voxels in which this global connectivity was significantly correlated to performance on the CFMT in both rest scans separately, with the ROIs from the face patch connectivity analysis overlaid. The peaks of the two analyses overlap almost entirely, except that the global connectivity analysis also picks out the face ROIs, which are absent in the previous analysis, while a few of the ROIs, most notably bilateral IFG, are missing in the global connectivity approach. We carried out a similar analysis looking at the correlation between the global connectivity and the CCMT scores but found no significant clusters.

**Figure 3.**
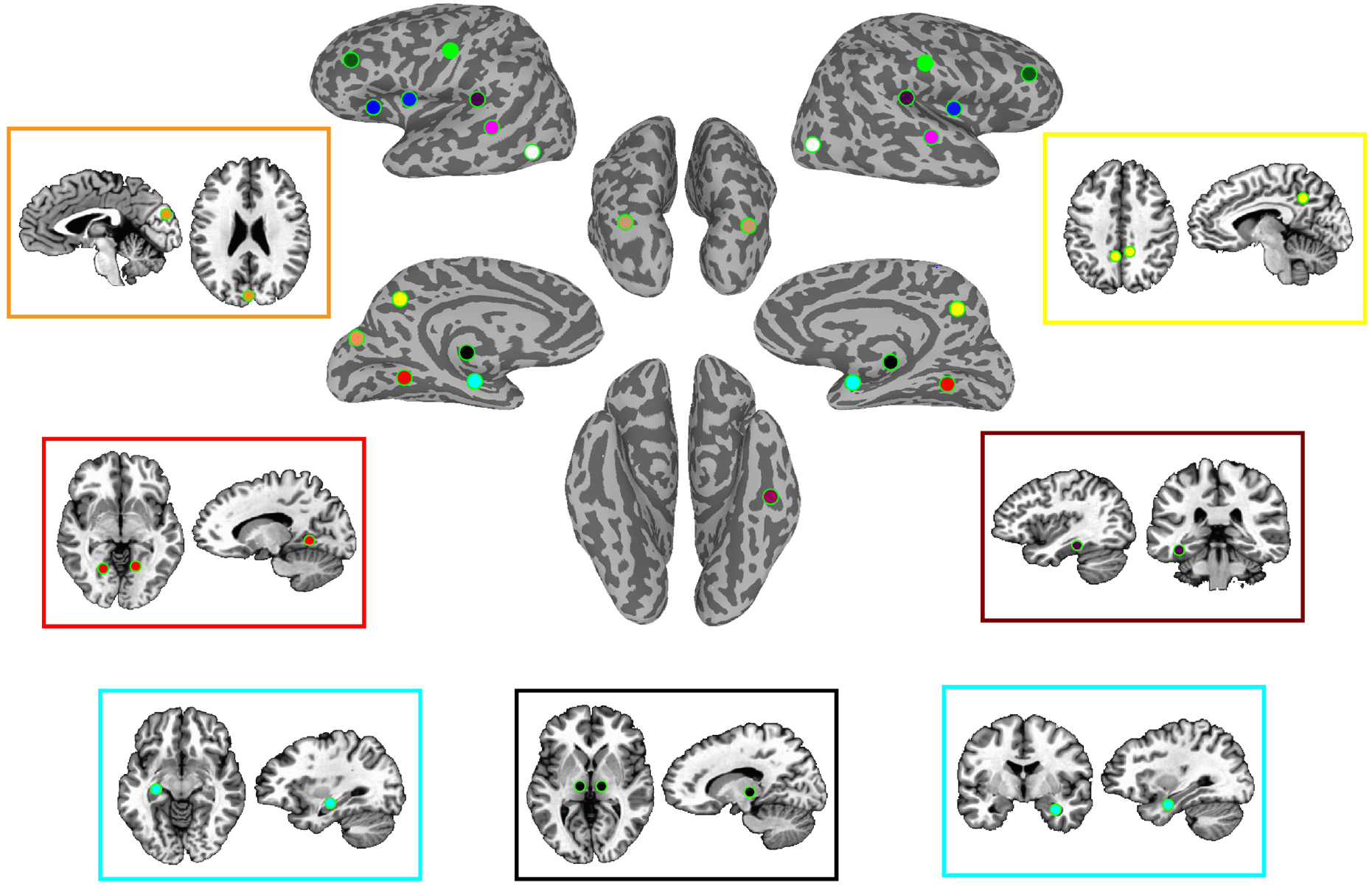
Group defined ROI locations. Locations of all 25 group defined ROIs. Light green: somatosensory, dark green: IFG, blue: insula and anterior insula, dark purple: STG/auditory cortex, magenta: STS, brown: dorsal attention stream, yellow: medial parietal, orange: cuneus, black: thalamus, cyan: hippocampus, red: parahippocampus, dark red: parahippocampus2.

### Examining the relationship among all nodes

To gain a better understanding of the network structures involving these ROIs, we calculated the full correlation matrix between all 32 ROIs, consisting of the seven face ROIs defined from the localizer, and the 25 ROIs added in the face connectivity analysis described above, averaged across all participants and both rest scans. The results are shown in Supplementary Figure 4. Predictably, correlations between homologous regions are the highest, as are correlations between FFA and OFA, and the insula with somatosensory cortex. To see which connections were most predictive of face memory abilities, as opposed to simply which areas were most strongly correlated, we again carried out the second order correlation analysis, calculating the correlation across participants between the correlations of each pair of ROIs, and the CFMT scores. This analysis was carried out for each rest scan separately, and we once again constrained the results by requiring that these correlations be significant across both rest scans.

The resulting second order correlation matrix, shown in Figure 5, shows all the ROI pairs for which the correlation between them was significantly predictive of performance on the CFMT, in both rest scans. Surprisingly, correlations within the face ROIs defined by the localizer (i.e. OFA, FFA, ATL, amygdala) were not significantly predictive of face memory abilities. The most predictive connections were between the face patches and medial temporal lobe structures as well as somatosensory cortex, within medial temporal lobe, and between STG/somatosensory cortex to the medial temporal lobe. To ensure that the predictive value of an ROI pair was not due to the degree of variance between participants in the first order correlation between the two nodes, we calculated the variance in the correlations across participants between each pair of ROIs and tested whether there was a correlation between this variance and the predictive value of the ROI pairs, but found none (r = 0.0015, p = 0.97).

**Figure 4.**
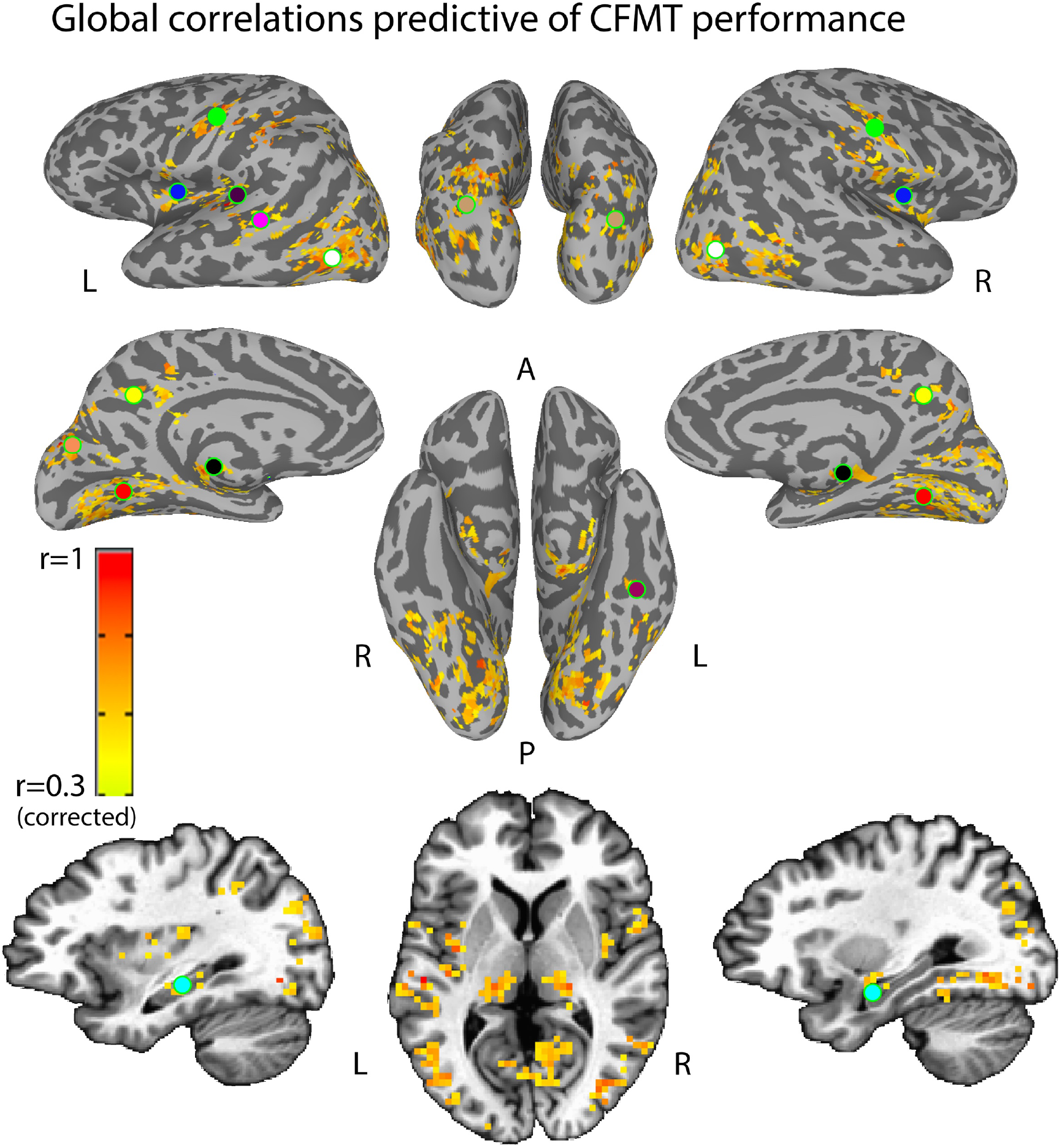
Global connectivity. Map shows voxels whose global connectivity, i.e. average connectivity with all the other voxels, during rest is significantly correlated to performance on the CFMT, after corrections for multiple comparisons. Overlaid are the ROIs defined from the previous analysis which was shown in Figure 3, using the face ROIs as seeds. Colors of the ROIs as in Figure 3.

**Figure 5.**
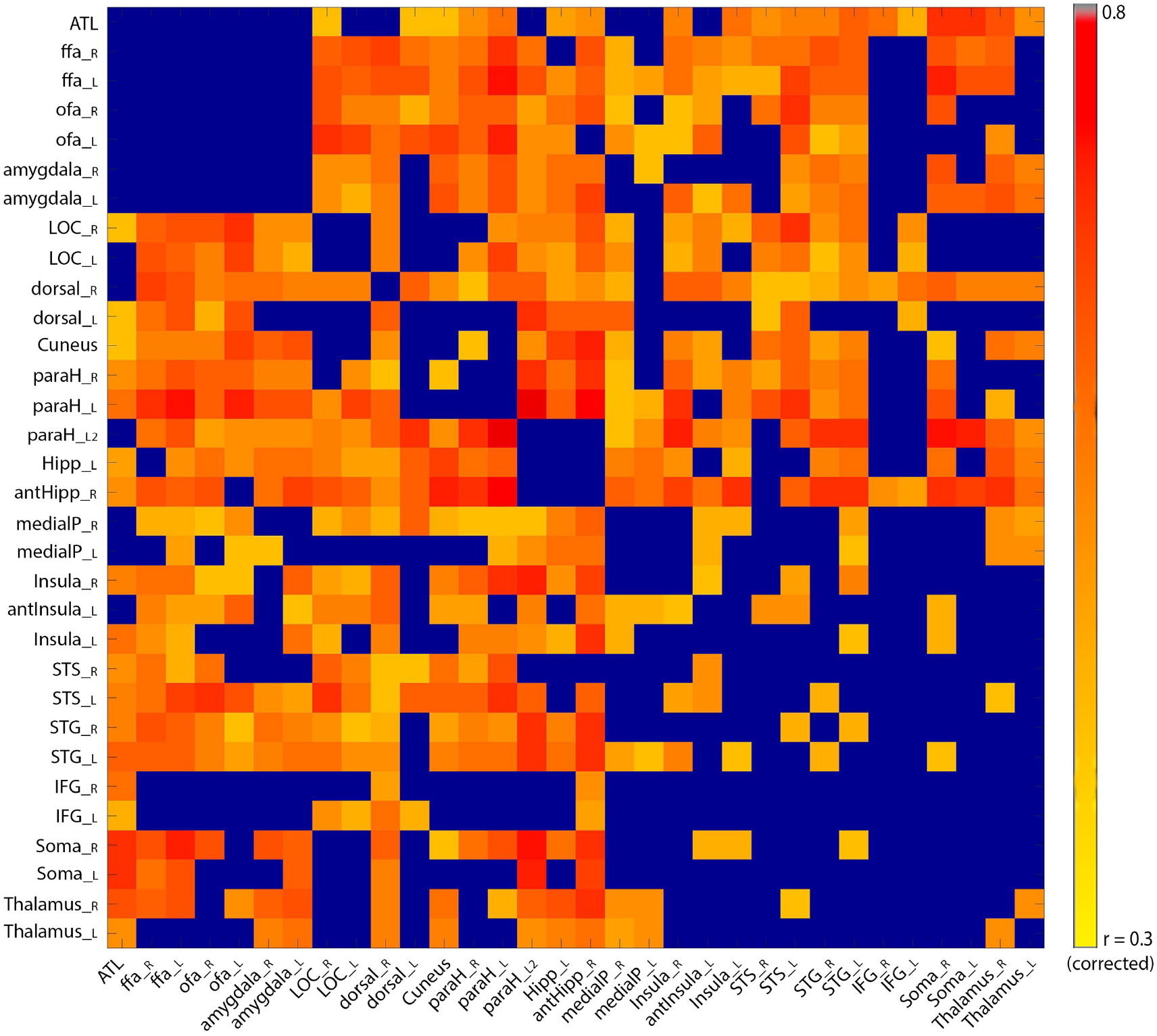
ROI pair correlations with behavior. Second order correlations of the correlation between each of the ROI pairs (consisting of the 7 individually localized ROIs from the localizer, and the 25 group-defined ROIs from the seed analysis) and CFMT. Values indicate how predictive the correlation between each ROI pair is of performance on the CFMT. Blue denotes an insignificant correlation. Note the lack of significant predictive value of the correlation between the different face ROIs (top left).

### Specificity for face memory

To further test whether the new ROIs, defined through their predictive value for the CFMT, are involved specifically in face memory or rather underpin more domain general memory processes, we redid the analysis of the second order correlation of the between ROI pair correlations, this time to the CCMT. The only ROI pair whose correlation was significantly predictive of scores on the CCMT, was right medial parietal with right LOC (r = 0.44, p = 0.0013). To more directly test the degree of variance explained by domain general rather than face specific processes in the predictive power of the connections within our network, we calculated the second order partial correlations of the correlation of each ROI pair with the CFMT scores, accounting for the CCMT scores, and then examined the difference for each ROI between the correlation with the CFMT, and the partial correlation. Supplementary Figure 5 shows this difference score, between the two correlation measures. We next ran a permutation test to determine the threshold at which this difference can be considered significant (see methods), and the only significant differences were found in the correlation of the right and left medial parietal regions to right LOC (correlation difference = 0.1 and 0.085, p~0.021, p~0.045 respectively).

### Comparing rest to movie viewing

Finally, we asked whether the network we uncovered could in some way be relevant only during rest, or whether the same network is also predictive of face memory abilities during a behaviorally pertinent task, such as naturalistic movie viewing of a scene involving faces. To examine this, we took the data from the movie viewing run, calculated the global connectivity for each voxel as above, and the second order correlation across participants of the global connectivity of each voxel with the CFMT scores. We then compared the predictive value of each voxel during rest, to its predictive value during the movie. These were highly correlated across voxels (r = 0.56, p = 0), as is shown in Figure 6.

**Figure 6.**
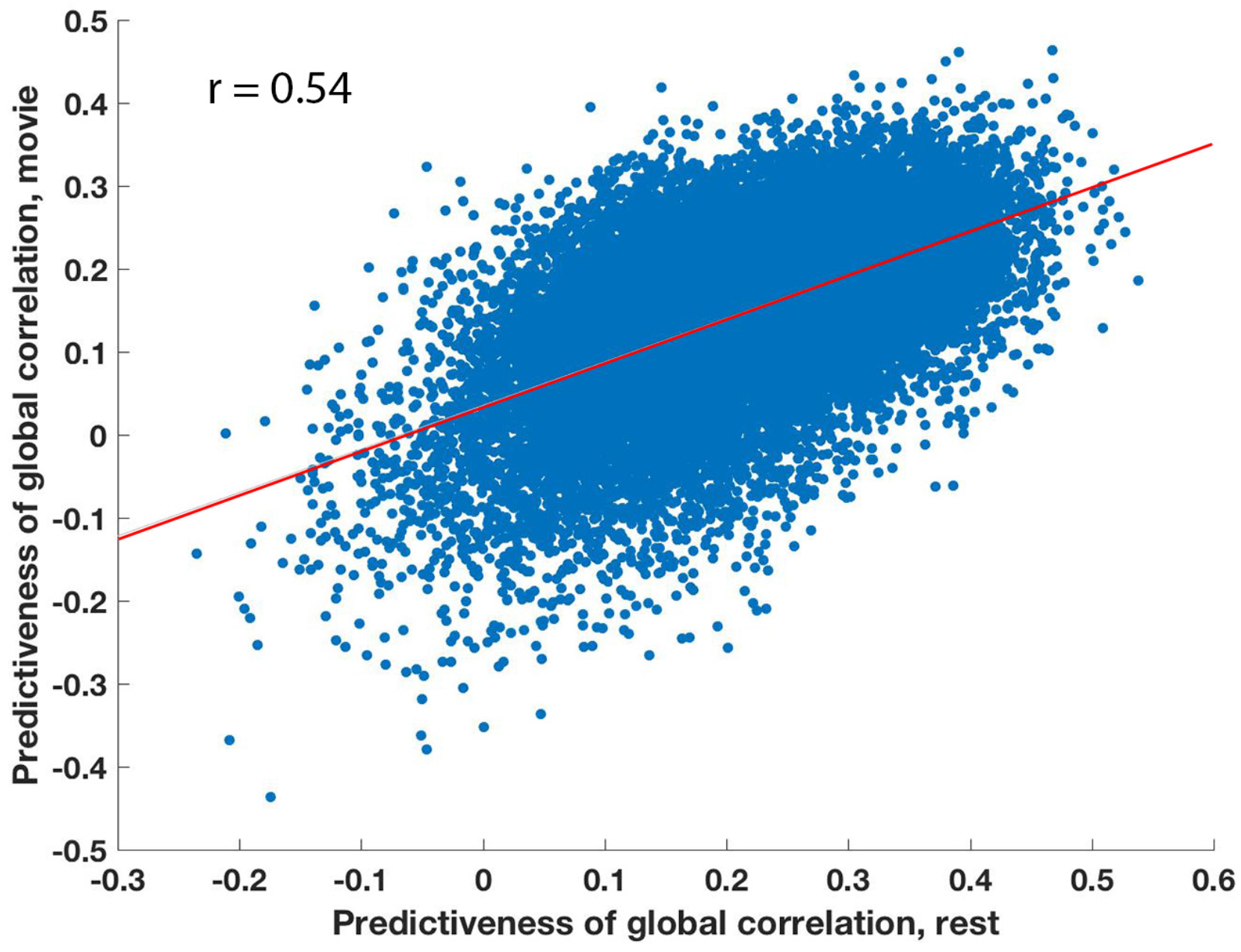
Comparison to movie data. Second order correlation of the global connectivity of each voxel with performance on the CFMT at rest (x-axis) and during the movie (y-axis). Predictiveness of the voxels across rest and movie viewing was highly correlate (r = 0.54), indicating that these networks are relevant for face memory during naturalistic viewing of faces, and not only during rest.

## Discussion

The focus of this current work was to identify additional networks beyond the traditional face network which are involved in face recognition memory. Using second order correlations to search for links between connectivity and behavior, we were able to uncover a number of regions whose connectivity either with the ventral face patches, or with each other, strongly and significantly predicted face memory abilities, as measured by the CFMT (Figures 3-4). Surprisingly, it was not correlations within the face regions which were most predictive, and in fact correlations between FFA/OFA/ATL and amygdala did not significantly predict face memory abilities (Figure 5, and note also the prominent absence of the other face patches in the seed based correlation maps shown in Supplementary Figures 1-3), although the degree of selective activation for faces within those regions was predictive (Figure 2). The combination of these two findings suggests that while these visual regions are clearly crucial for face memory, as evidenced by the link to selective activation for faces, the information transfer between them is not what drives face memory abilities. Rather, looking at the whole brain analysis, we find evidence that memory for faces, even unknown faces such as those presented in the CFMT, extends well beyond the visual face patches, and is supported by widespread networks involved in memory, social cognition, and even auditory processing.

The predictive peaks that came up in our analysis can be roughly divided as belonging to memory related, social, visual/attention and auditory regions. Medial temporal lobe regions, such as the hippocampus and parahippocampus, have long been associated with memory processes ^25, 26^, as have medial parietal regions ^27, 28^.On the other hand, somatosensory cortex has been found in multiple studies to be implicated in social processing ^29-31^, and this same region of somatosensory cortex identified here, was previously found to be under-connected both globally, and specifically to other social brain regions such as STS, in patients with Autism Spectrum Disorder compared with typically developing individuals ^32-34^. Insular cortex has been shown to receive input from somatosensory cortex among others ^35^, and to also be involved in social/emotional processing ^36, 37^.One of the most unexpected findings was that face memory performance was strongly predicted by the correlation between the peaks found in STG, along Heschel’s gyrus ^38-40^ and regions of the face processing network, (in particular right FFA; r = 0. 45). This is congruent with previous findings linking listening to voices to FFA activation, even in the absence of face stimuli ^41, 42^.

The same key predictive regions were also found using the data driven global connectivity approach, with the most obvious difference between the seed based analysis and global connectivity analysis being the presence of peaks in FFA, OFA and amygdala in the latter. These face regions were absent from the seed based analysis, as the correlation between the face regions is not predictive of performance on the CFMT. However, their correlations with many other regions are relevant to behavior (as can be seen in Figure 5), which is why they come to light in the global connectivity analysis. There was an additional peak found in this analysis centered around ATL, but it did not survive the cluster size correction for multiple comparisons.

The other intriguing finding relates to the specificity of these connections for face memory. The Cambridge Car Memory Test is identical to the Cambridge Face Memory Test in format and is matched for difficulty ^24^, and therefore involves the same general cognitive and memory processes, with the only difference being the object of memory, cars in one and faces in the other. That there were no significant second order correlations elsewhere in the brain to the CCMT with the face patches is expected, as the face patches were defined specifically by their selectivity for faces, and it is therefore unsurprising that they are not involved in memory for other visual objects such as cars. However, the 25 new ROIs which were defined could equally underlie domain general memory processes rather than face specific ones, and yet the only ROI pair which significantly predicted performance on the CCMT was the right medial parietal ROI, with right LOC. Similarly, when regressing out the variance explained by the CCMT, the predictive value of most ROI pairs to the CFMT did not significantly change (Supplementary Figure 5), with the only exception again being the medial parietal ROIs with right LOC. Without an individual car/object localizer it is difficult to directly compare the predictive value of each of these ROIs separately for face memory vs. car memory, but from the above partial correlation analysis it appears that the network described in these results, with connections not only between the face patches and memory/social regions, but also within memory/social regions, is largely specialized specifically for face memory (with the possible exception of the medial parietal regions).

The data driven approach, using the global connectivity of each voxel as an input to the second order correlations with performance on the CCMT, also failed to find any significant clusters whose connectivity predicts the CCMT scores. This approach is limited in that it is biased toward regions with widespread task-relevant connectivity. Regions with limited relevant connectivity will not be identified by this method, as was the case for IFG in correlation with the CFMT scores, as IFG connects to only a few other ROIs in a way that is relevant for CFMT performance. It is therefore possible that there are other regions which subserve memory for other types of objects, but this network is more spatially constrained than the one for faces. Seed regions identified through a specific localizer for cars/objects might have uncovered regions which underpin memory for objects.

Correlations observed during resting state scans are generally interpreted as representing the baseline of the brain, echoing meaningful structures which exist during task performance ^43, 44^.It is therefore interesting to compare the results from the resting state scans to the results during the movie, where participants were viewing actual faces in a naturalistic setting. Apart from adding an additional independent dataset, the high correspondence between the predictive value across voxels at rest and during the movie helps fortify the claim that these networks have real-world relevance for face memory.

Taken together, these findings suggest a more holistic identity processing underlying face memory, which is more widespread than memory processing for cars. The networks underlying face memory seem to integrate visual information with social and auditory cues, perhaps taking into account elements such as voice, and the emotions elicited by particular faces. That this same network underlies unknown faces such as those in the CFMT, is an indicator of the framework in which we process a novel face. Some of the memory regions identified in this study, such as the region in hippocampus as well as the parahippocampal regions, appear to be strongly biased toward face processing if not specific to faces, and warrant further study to determine the degree of their selectivity.

## Methods

### Participants

Fifty three participants (24 female) aged 16 – 30 (mean age = 23.1) were recruited for this experiment. All participants were screened for any history of neurological and psychiatric disorders. In addition, all participants had normal to corrected to normal vision. One participant was excluded from analysis because of abnormal brain structure, and two were excluded due to inattention on behavioral testing. The experiment was approved by the NIMH Institutional Review Board (protocol 10-M-0027). Written informed consent was obtained from all participants.

### Behavioral testing

Prior to the scan, all participants completed two memory tasks: the Cambridge Face Memory Task (CFMT) ^23^ and the Cambridge Car Memory Test (CCMT) ^24^. All but 4 subjects completed the memory tasks directly before the scan. The CFMT is comprised of three parts; in the first part, participants are shown three views of a target face, and then presented with a forced-choice test with the target face and two distractor faces. Participants had to select the face that matched the original target face. There are six target faces, each of which was presented three times, for a total for eighteen trials. In the second part, participants were presented with frontal views of the six target faces for 20 seconds, followed by 30 forced-choice tests with one target face and two distractor faces. Next, subjects were presented with the frontal views of the six target faces for 20 seconds, followed by 24 more forced-choice test displays presented with a Gaussian noise overlay. The CCMT uses the same structure as the CFMT, but uses cars, instead of faces. For both the CFMT and the CCMT, recognition scores were the sum correct responses on the three sections. Subjects who had two or more incorrect trials ln the first introductory phase of the memory tests were excluded due to concerns about attention to tasks.

### Imaging data collection and MRI parameters

All scans were performed at the Functional Magnetic Resonance Imaging Core Facility on a 32 channel coil GE 3T (GE MR 750 3.0T) magnet and receive-only head coil, with online slice time correction. The scans included a 6 minute T1-weighted magnetization prepared rapid gradient echo (MPRAGE) sequence for anatomical co-registration, which had the following parameters: TE = 2.7, Flip Angle = 12, Bandwidth = 244.141, FOV = 30 (256 x 256), Slice Thickness = 1.2, axial slices. Functional images were collected using multi-echo acquisition using the following parameters: TR = 2s, voxel size = 3*3*3, flip angle = 60, multi-echo slice acquisition with 3 echos, TE = 17.5ms, 35.3ms, and 53.1ms, Matrix = 72×72, slices = 28. 270 TRs were collected for the rest scans, 250 TRs for the face/scene localizer scans, and 285 TRs for the movie. All scans used an accelerated acquisition (GE’s ASSET) with a factor of 2 in order to prevent gradient overheating.

### Scan stimuli and task

Each scan started with two 9 minute rest scans. During this scan, participants were presented with a uniformly grey screen with a fixation cross. Participants were instructed to lie still, not fall asleep and look at the screen. After the rest scans, participants completed two runs of an 8 minute and 20 second face/scene localizer scan. Four subjects completed a different, 9 minute 20 second localizer scan, and they were excluded from the face selectivity analysis relying on beta weights, as those were not comparable between localizer types. Each localizer began with a 20 second blank grey screen, followed by sixteen 20 second presentation blocks and a 10 second blank grey screen with a fixation cross. During presentation blocks, 20 pictures of faces (face blocks) or scenes (scene blocks) were presented (stimulus duration = 200ms, interstimulus interval = 700ms), with one or two images repeating in succession in each block. Subjects were instructed to look for these repetitions (1-back task) and respond using a button box. There were 8 face blocks and 8 scene blocks in each localizer run, with 320 exemplars from each category. Each exemplar repeated no more than twice in each run. After the two localizer runs, all subjects were shown a clip from *The Princess Bride*.This video started with 30 seconds of uniformly grey screen with a fixation cross at the center, followed by a 9 minute clip from the movie. For the movie, subjects were instructed to stay still, keep their eyes open and watch the video.

### MRI offline data pre-processing

Post-hoc signal pre-processing for the functional images was performed in AFNI ^45^. The first four EPI volumes from each run were removed to ensure all volumes were at magnetization steady state. Any large transients that remained were removed using a squashing function (AFNI’s 3dDespike). Volumes were slice-time corrected and motion parameters were estimated with rigid body transformations (using AFNI’s 3dVolreg function). Volumes were co-registered to the anatomical scan. The data were then processed using AFNI’s meica.py to perform a multi-echo ICA analysis (ME-ICA). This process removes nuisance signals such as hardware-induced artifacts, physiological artifacts and residual head motion ^46^.

### ROI Selection

The localizer data was used to define individual face ROIs for each subject. A standard General Linear Model was used with a 20 second long boxcar function. This was convolved with a canonical hemodynamic response function, and deconvolved using the AFNI function 3dDeconvolve. Face selective ROIs were found using the faces>scenes contrast. The functional and anatomical datasets were co-registered using AFNI, then transformed to Talaraich space.All ROIs for each individual participant, were defined in Talaraich space. In the faces>scenes contrast, we identified the center of mass for the bilateral fusiform face area (FFA), occipital face area (OFA) and amygdala, in addition to the right anterior temporal lobe (ATL) face patch. We then defined a spherical ROI of 6mm radius around each of these centers of mass to obtain 7 individually localized visual ROIs.

### Whole brain analysis cluster size correction

In addition to these individually localized ROIs, we obtained 25 group-defined ROIs, by using each of the seven individually defined ROIs as a seed, and calculating for each voxel the second order correlation across participants between that voxel’s correlation with the seed for each participant (transformed to z-scores using Fisher’s transform), and the CFMT / CCMT scores, as appropriate. We did this for each of the rest scans separately, and then combined the resulting maps for each seed across both rest scans, by requiring that voxels be significantly correlated with behavior in both rest scans at either p~0.05 or p ~ 0.01 in order to be counted, resulting in seven maps, one per seed. We then ran a permutation test cluster size correction for multiple comparisons, for all seven maps together, by permuting the CFMT scores 10,000 times and then testing second order correlations for each voxel, for each seed, and requiring that voxels be significantly correlated in both rest scans at either p~0.05 or p ~ 0.01 to be counted. We then took the largest cluster at the 95% percentile across all maps as our minimum cluster size for p~0.05 and p~0.01. We identified peaks in the surviving clusters across the seven seed maps at either significance threshold, and defined new ROIs as 6mm spheres around the peaks, resulting in 25 ROIs for CFMT, and none for CCMT. The 7 individually localized visual ROIs, in addition to the 25 group-defined ROIs, were used as targets in subsequent analyses. Minimum cluster size for the global connectivity analysis was determined in the same way.

## Data Analysis

All data were analyzed with in-house software written in Matlab, as well as the AFNI software package. Data on the cortical surface were visualized with SUMA (SUrface MApping) ^47^. Two-tail t-tests were used for all p-values on correlations, unless otherwise stated. For the permutation test used to determine the threshold of significance for the difference between the second order Pearson’s correlation of each ROI pair’s correlation (per participant) with the CFMT scores, to the same second order correlation but with the CCMT scores as a regressor, the correlation and partial correlation (calculated used MATLAB’s partialcorr function) scores for each ROI pair were first converted to z-scores using Fisher’s transform, and the difference between them was calculated. For each ROI pair, for 10,000 iterations, we then permuted the subject labels on the correlation values between that ROI pair, and calculated the permuted second order correlation to the CFMT, the second order partial correlation to the CFMT with the CCMT scores regressed, and the difference between them, for each rest scan. We then took the average for each iteration across the two rest scans, and set as the threshold the result of the 95^th^ percentile across all iteration across all possible ROI pairs. Supplemtary Figure 5 shows the difference between the correlation to the partial correlation for all ROI pairs, averaged across the two rest scans, though only two ROI pairs showed a significant difference in scores.

## Acknowledgments

We thank Adrian Gilmore, Cibu Thomas and Stephen Gotts for helpful conversations and insights. This work was supported by the Intramural Research Program, National Institute of Mental Health (ZIAMH002920), clinical trials number NCT01031407.

---

**Supplementary Figure 1.**
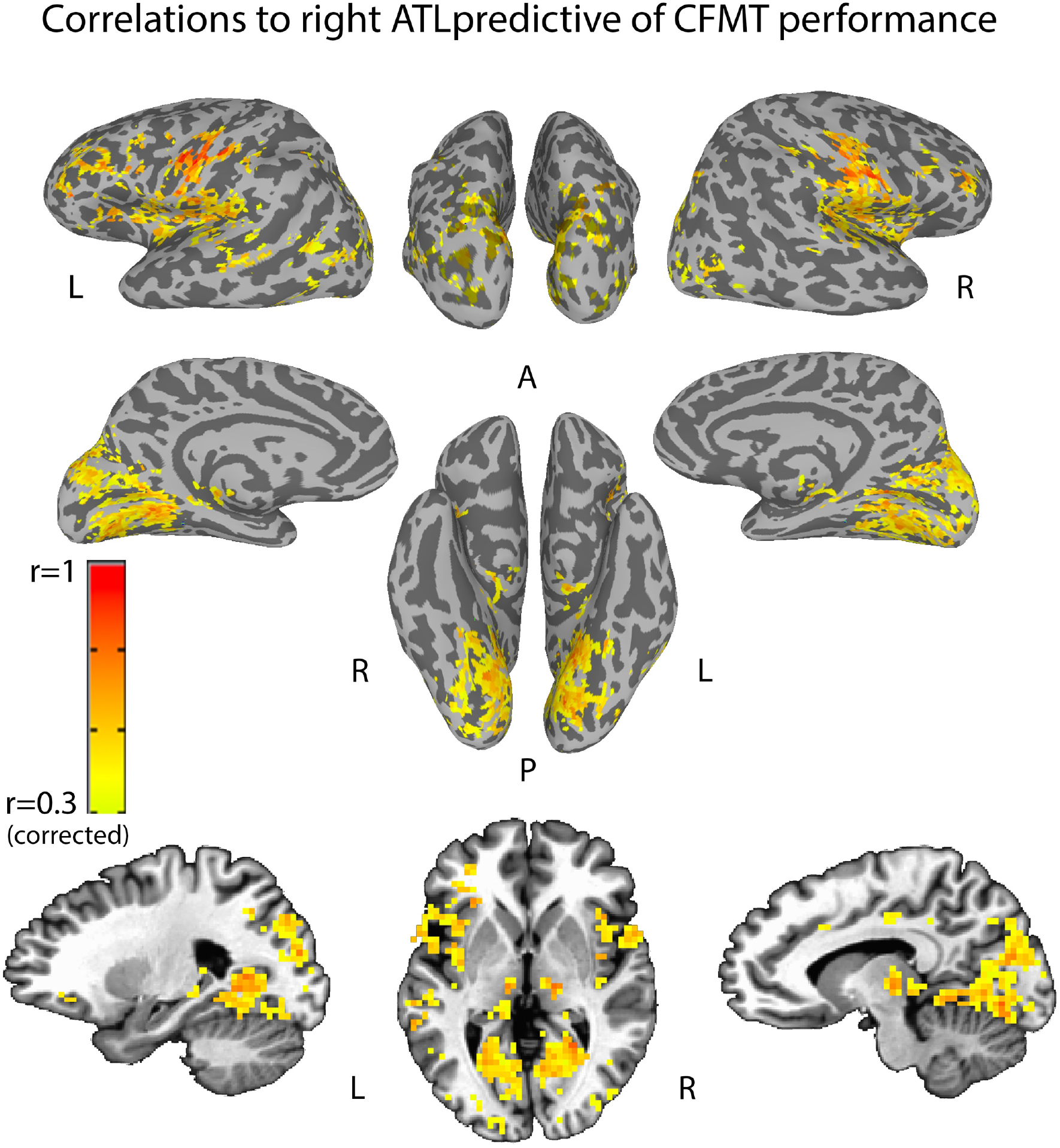
Clusters showing voxels in which there was a significant second order correlation between their correlation to the ATL seed and performance on the CFMT, indicating that the correlation with ATL is predictive of behavior, in both rest scans separately. Maps corrected for multiple comparisons through a cluster permutation test analysis (see methods). Note large clusters in the parahippocampus and along the occipital part of the parieto-occipital sulcus, in somatosensory cortex, IFG, insula, STG, STS and along the dorsal visual attention stream.

**Supplementary Figure 2.**
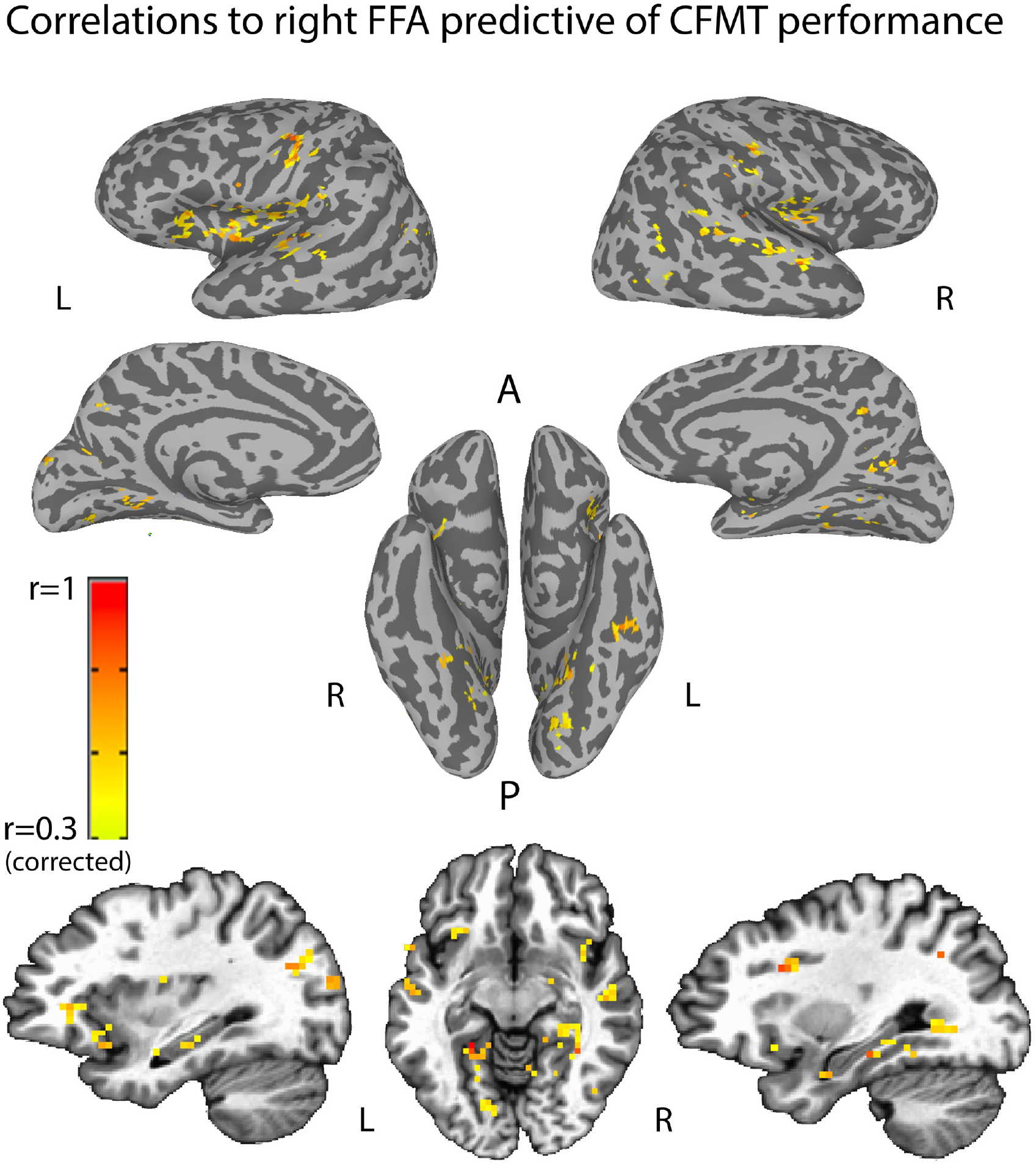
Clusters showing voxels in which there was a significant second order correlation between their correlation to the right FFA seed and performance on the CFMT, indicating that the correlation with right FFA is predictive of behavior, in both rest scans separately. Maps corrected for multiple comparisons through a cluster permutation test analysis as above (see methods). Note peaks in the parahippocampus, in somatosensory cortex, insula, STG, STS, medial parietal cortex and hippocampus.

**Supplementary Figure 3.**
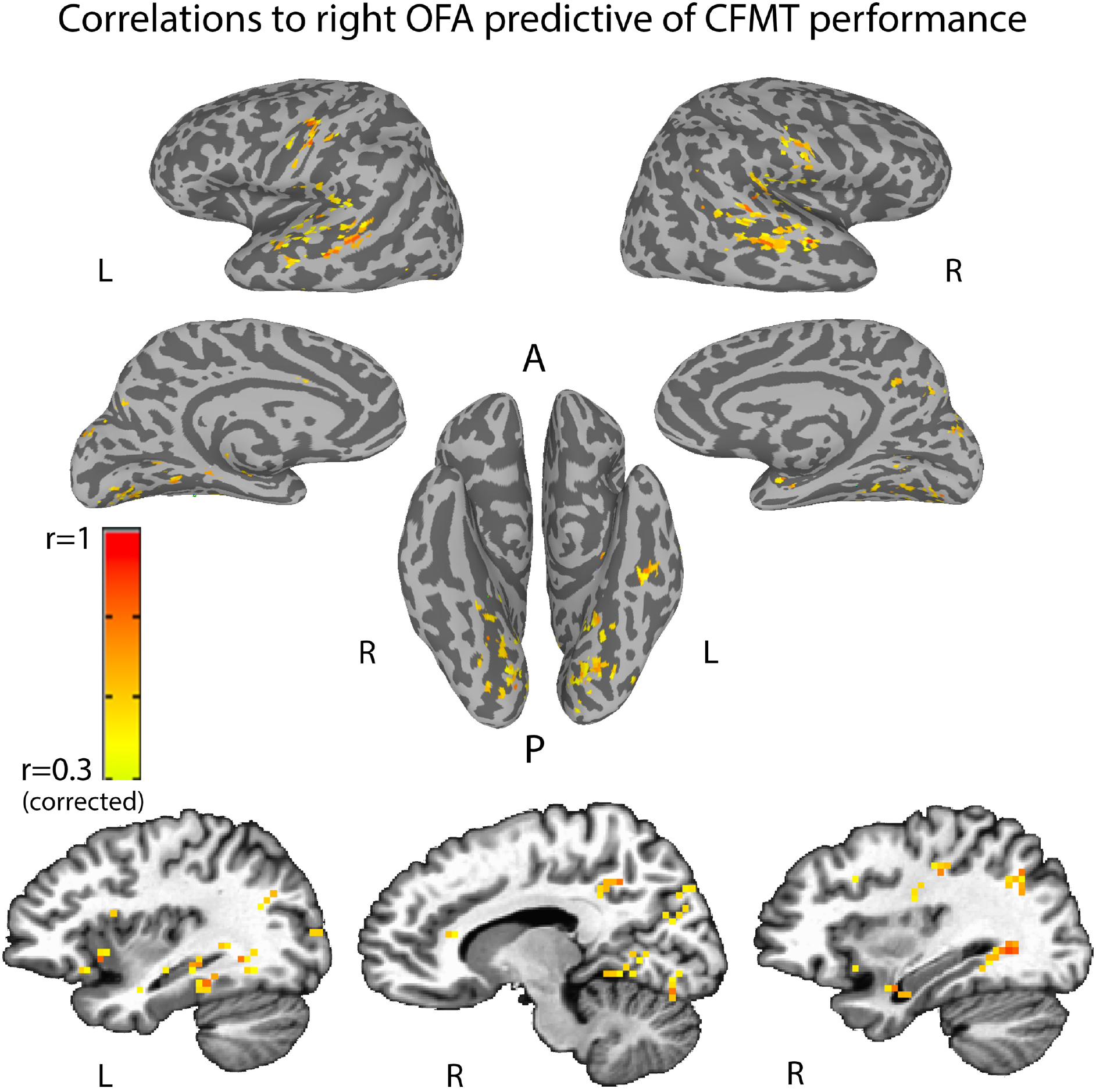
Clusters showing voxels in which there was a significant second order correlation between their correlation to the right OFA seed and performance on the CFMT, indicating that the correlation with right OFA is predictive of behavior, in both rest scans separately. Maps corrected for multiple comparisons through a cluster permutation test analysis as above (see methods). Note peaks in the parahippocampus, in somatosensory cortex, STG, STS, medial parietal cortex and hippocampus.

**Supplementary Figure 4.**
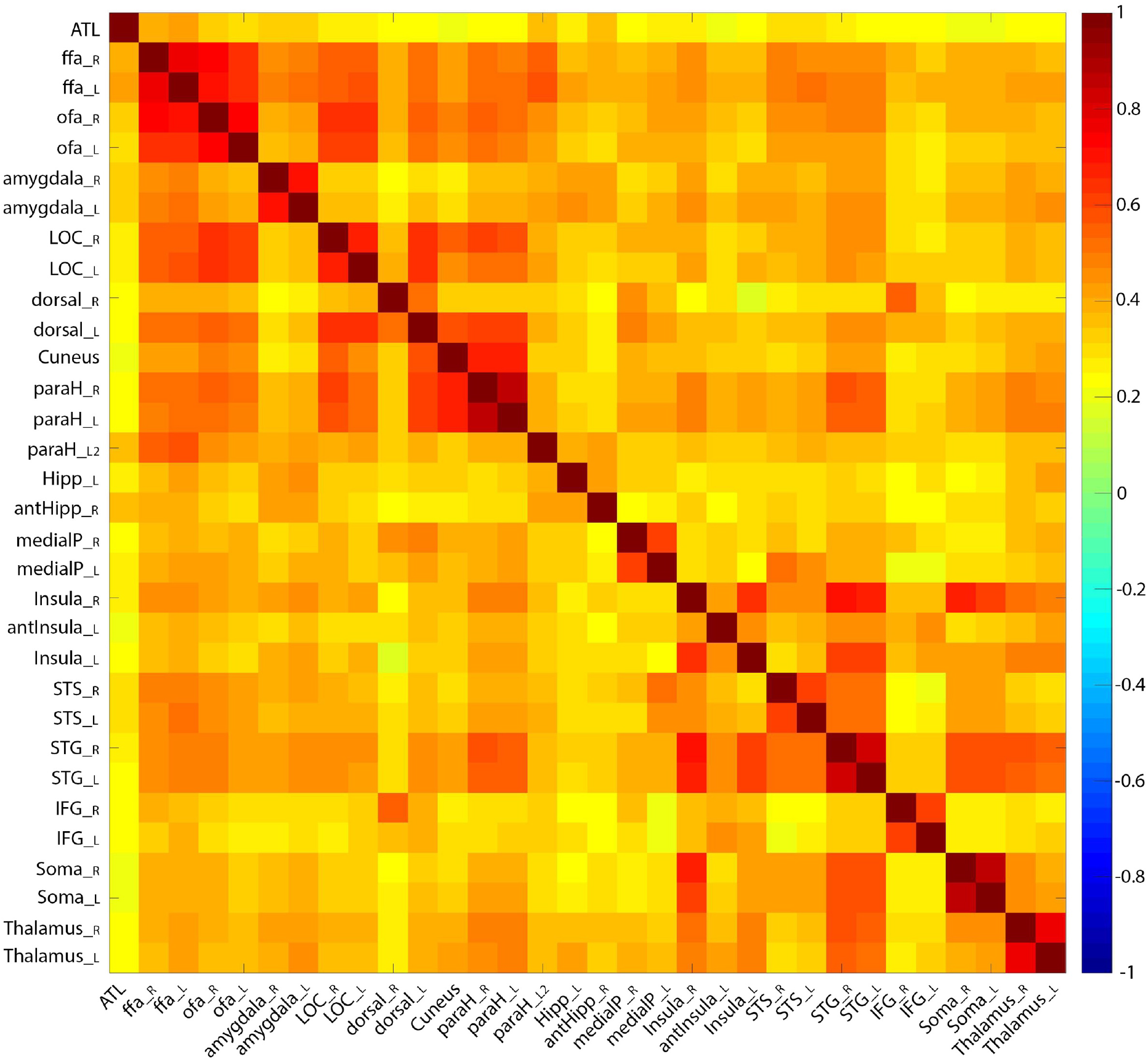
First order correlations between all 32 pairs of ROIs identified either through the localizer, or through the second order correlation seed analysis. Correlations between homologous regions are strongest, as expected, as are correlations between FFA and OFA and LOC.

**Supplementary Figure 5.**
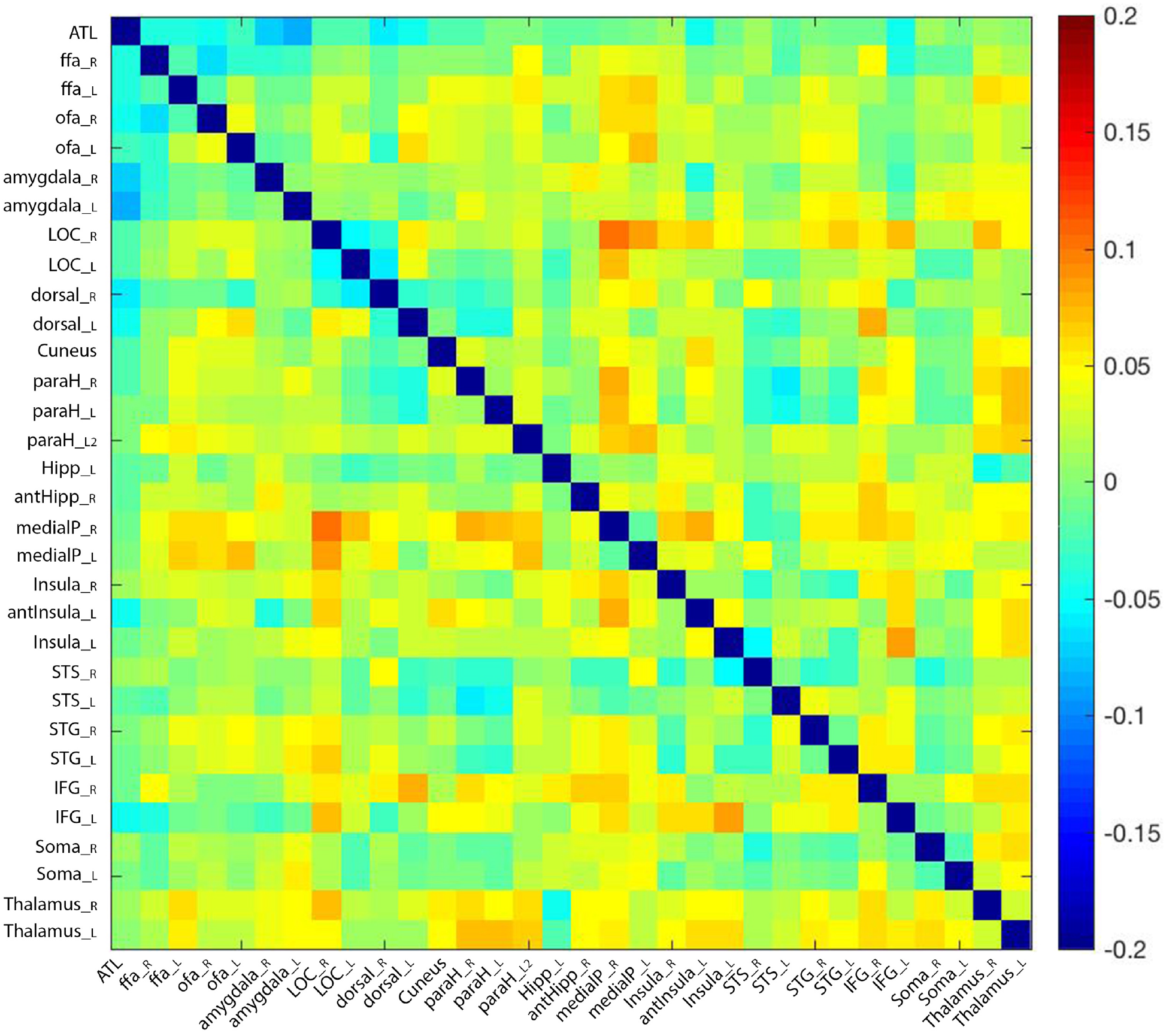
The difference between second order correlation of correlations between each ROI pair with CFMT scores, and the partial correlation of that ROI pair with CFMT while controlling for CCMT scores, averaged across both rest runs. The only significant difference was found between the right and left medial parietal ROIs, to right LOC.

